# Neural crest and periderm-specific requirements of *Irf6* during neural tube and craniofacial development

**DOI:** 10.1101/2024.06.11.598425

**Authors:** Shannon H. Carroll, Sogand Schafer, Eileen Dalessandro, Thach-Vu Ho, Yang Chai, Eric C. Liao

**Affiliations:** Center for Craniofacial Innovation, Children’s Hospital of Philadelphia Research Institute, Children’s Hospital of Philadelphia, PA 19104, USA; Center for Craniofacial Molecular Biology, University of Southern California, Los Angeles, CA USA; Division of Plastic and Reconstructive Surgery, Department of Surgery, Children’s Hospital of Philadelphia, PA 19104, USA; Shriners Hospital for Children, Tampa, FL 33607, USA

**Keywords:** Irf6, cleft palate, neural tube, neural crest, Van der Woude Syndrome, periderm

## Abstract

*IRF6* is a key genetic determinant of syndromic and non-syndromic cleft lip and palate. The ability to interrogate post-embryonic requirements of *Irf6* has been hindered, as global *Irf6* ablation in the mouse causes neonatal lethality. Prior work analyzing *Irf6* in mouse models defined its role in the embryonic surface epithelium and periderm where it is required to regulate cell proliferation and differentiation. Several reports have also described *Irf6* gene expression in other cell types, such as muscle, and neuroectoderm. However, analysis of a functional role in non-epithelial cell lineages has been incomplete due to the severity and lethality of the *Irf6* knockout model and the paucity of work with a conditional *Irf6* allele. Here we describe the generation and characterization of a new *Irf6* floxed mouse model and analysis of *Irf6* ablation in periderm and neural crest lineages. This work found that loss of *Irf6* in periderm recapitulates a mild *Irf6* null phenotype, suggesting that *Irf6*-mediated signaling in periderm plays a crucial role in regulating embryonic development. Further, conditional ablation of *Irf6* in neural crest cells resulted in an anterior neural tube defect of variable penetrance. The generation of this conditional *Irf6* allele allows for new insights into craniofacial development and new exploration into the post-natal role of *Irf6*.

## Introduction

*IRF6* was one of the first genetic determinants of syndromic cleft lip and palate malformation, uncovered from genome-wide association studies of Van der Woude syndrome (VWS) and popliteal pterygium syndrome (PPS) (Kondo et al., 2002). *IRF6* gene variants are also major contributors to non-syndromic cleft lip with or without cleft palate (Leslie et al., 2013; Park et al., 2007; Rahimov et al., 2008; Zucchero et al., 2004). Multiple studies using mouse and zebrafish models have shown that *Irf6* is expressed in the basal epithelium and periderm during embryonic development with dynamic expression in the oral epithelium during palatogenesis (Carroll et al., 2020; de la Garza et al., 2013; Dougherty et al., 2013; Ferretti et al., 2011; Ingraham et al., 2006; Iwata et al., 2013; Knight et al., 2006; Kousa et al., 2017; Richardson et al., 2006; Xu et al., 2006). *Irf6* is necessary for keratinocyte differentiation (Biggs et al., 2012; Ingraham et al., 2006; Restivo et al., 2011; Richardson et al., 2006) and for the development of the periderm (de la Garza et al., 2013; Li et al., 2017; Richardson et al., 2009; Richardson et al., 2014; Sabel et al., 2009). Ablation of *Irf6* in mice resulted in severe epithelial adhesions that caused “cocooning” of the embryo and caused adherence of palatal shelves to the tongue in the vertical orientation precluding elevation and fusion of the secondary palate (Ingraham et al., 2006; Richardson et al., 2009; Richardson et al., 2014) and fuse (Iwata et al., 2013).

While most studies have examined the requirement of *Irf6* in epithelial differentiation, several studies have described *Irf6* function in non-epithelial tissue, in either autonomous or non-cell-autonomous fashion (Goudy et al., 2013; Thompson et al., 2019). *Irf6* is expressed in cell types not restricted to surface epithelium during early development, including the craniofacial mesenchyme and neuroectoderm (Carroll et al., 2020; Fakhouri et al., 2017; Goudy et al., 2013; Sabel et al., 2009; Thompson et al., 2019). Further, analysis of murine *MCS9.7* enhancer element activity, which replicates endogenous *Irf6* expression in most tissues, yielded expression in developing somites, tongue, axial cartilage, and muscle (Fakhouri et al., 2012).

We and others have described that mesenchymal-derived craniofacial tissue, such as muscle and cartilage are dysmorphic in the *Irf6* null mice (Carroll et al., 2020; Chu et al., 2016; Thompson et al., 2019). However, it remains unclear whether there is a cell-autonomous role of *Irf6* in non-epithelial cell types, a non-cell-autonomous role caused by loss of epithelial *Irf6*, or if these dysmorphologies are associated consequences of the severe epithelial adhesions caused by a dysfunctional epithelium.

In addition to orofacial and epithelial development, *Irf6* has been found to have a role in neurulation. *Irf6* is expressed in the neuroectoderm of the neural folds and is co-expressed with *Tfap2a,* a known regulator of neural tube closure (Kousa et al., 2019). Although neural tube defects are not apparent in *Irf6* null mice, ablation of other genes in the Irf6 regulatory pathway, i.e. *Tfap2A* and *Grhl3*, leads to rostral and caudal neural tube defects (Schorle et al., 1996; Ting et al., 2003; Zhang et al., 1996). Utilizing an *Irf6* hypomorph allele and a *Krt4:Irf6* transgenic mouse to titrate *Irf6* expression levels, Kousa et al. found homeostasis of *Irf6* to be required for neurulation (Kousa et al., 2019).

The ability to interrogate non-epithelial and post-natal functions of *Irf6* has been impaired by the severe and lethal phenotype of the *Irf6* null mouse models. A previously generated *Irf6* floxed mouse model has given some insight (Smith et al., 2017). Conditional ablation of *Irf6* in oral epithelium via a *Pitx2*-Cre driver line resulted in tooth development and maturation defects (Chu et al., 2016). Since the previously generated *Irf6* floxed allele was reported to show variable recombination efficiency and we remained unsuccessful in acquiring it (Smith et al., 2017), we generated a new conditional *Irf6* floxed mouse allele for this work. This *Irf6* conditional allele demonstrated complete recombination efficiency with every ubiquitous and tissue-restricted Cre drivers we have tested.

In this study, we describe the generation of a new conditional *Irf6* mouse allele and analyze *Wnt1-*Cre2-mediated disruption of *Irf6* in the neural crest cells (NCCs). We also utilized the *Krt6ai*-Cre driver line to ablate *Irf6* function in periderm. These results demonstrate for the first time a cell-autonomous role for Irf6 in the neural crest as well as corroborate the functional role of Irf6 in the periderm during orofacial development.

## Materials and Methods

### Generation of a new conditional *Irf6* mouse allele

All procedures were approved by IACUCs for Massachusetts General Hospital and Harvard University where the initial work was carried out. The *Easi*-CRISPR protocol was utilized to introduce loxP sites (Miura et al., 2018) flanking exons 3 and 4 of *Irf6*. As these exons contain the DNA binding region (Kondo et al., 2002), they are predicted to be required for Irf6 transcriptional function and have been previously targeted for conditional ablation of Irf6 (Smith et al., 2017). Guide RNAs (gRNA) were designed within the intronic regions flanking exons 3 and 4 using the CRISPR gRNA design tool from Benchling and were ordered from Synthego. Single-stranded DNA (ssDNA) donor sequences were designed to contain the loxp sequence flanked by homologous arms and were ordered from IDT. Cas9, gRNA and donor ssDNA were injected into mouse zygotes by the Harvard Genome Modification Facility. Resulting viable pups were genotyped by PCR as well as sequenced to ensure the insertion of the loxP sequences within the same DNA strand. A female mouse was identified with the correct genome modifications, was phenotypically normal, and was designated F0. Breeding with a wildtype C57BL/6J mouse generated F1s, which were in-crossed to generate mice homozygous for the floxed *Irf6* allele (*Irf6*^fl/fl^).

### Mouse lines

To validate efficient Cre recombination and to confirm recombination ablates *Irf6* function, *Irf6*^fl/fl^ mice were bred to the Cre deleter lines CMV-Cre (Jackson Labs stock# 006054) and EIIa-Cre (Jackson Labs stock# 003724). The resulting pups (viable and non-viable) were phenotyped and genotyped. *Wnt1*Cre2 and *Sox10*Cre were obtained from Jackson Labs (stock# 022501 and 025807, respectively). *Krt6ai*-Cre came from Vesa Kaartinen. Crect line came from Russ Carstens but originated from Trevor Williams. For timed pregnancies, E0.5 was determined upon observation of a copulatory plug.

### Histology and *in situ* hybridization

Mice were fixed with 4% formaldehyde followed by cryoprotection in 15 and 30% sucrose. Tissues were embedded in OCT and 10 μm sections were made. Hematoxylin and Eosin staining was performed according to a standard protocol (Fischer et al., 2008) and slides were imaged with a Leica DM6 upright microscope and LAS X software.

RNAscope probes for mouse *Irf6*, *Wnt1,* and *Sox10* were designed and manufactured by Advanced Cell Diagnostics. RNAscope *in situ* hybridization was performed according to the manufacturer’s protocol (Advanced Cell Diagnostics). Slides were imaged using a confocal laser scanning microscope (Leica SP8) and image processing was performed using ImageJ version 2.0 (2018).

### MicroCT analysis and measurements

Scans were performed using a µCT40 benchtop scanner (Scanco Medical AG, Brüttisellen, Switzerland). Scans were acquired with a 15 µm^3^ isotropic voxel size, 70 kVP peak x-ray tube potential, 114 mA intensity, and 300 ms integration time. Morphometric landmarks were chosen as previously described (Ho et al., 2015) and measurements were made using Avizo software.

## Results

### *Irf6* is expressed in *Wnt1*+ neuroectoderm and neural crest cell-derived cranial mesenchyme

*Irf6* null mice exhibit a foreshortened midface as well as malformation of neural crest-derived maxilla (Fakhouri et al., 2017; Richardson et al., 2006; Thompson et al., 2019). *Irf6* gene dosage has also been found to impact neural tube closure (Kousa et al., 2019). It was previously reported that *Irf6* is expressed in the neuroectoderm and neural folds of mouse embryos (Bertol et al., 2022; Kousa et al., 2019). To examine whether cranial neural crest cells express *Irf6*, we analyzed *Irf6* mRNA expression by RNAscope *in situ* hybridization in mouse embryos during early craniofacial development. At E8 and E9, we found *Irf6* mRNA co-expressed with *Wnt1* in the neuroectoderm. *Irf6* mRNA expression was also found co-expressed with *Sox10*, demonstrating *Irf6* expression in migratory neural crest cells (Fig 1b).

**Fig. 1.**
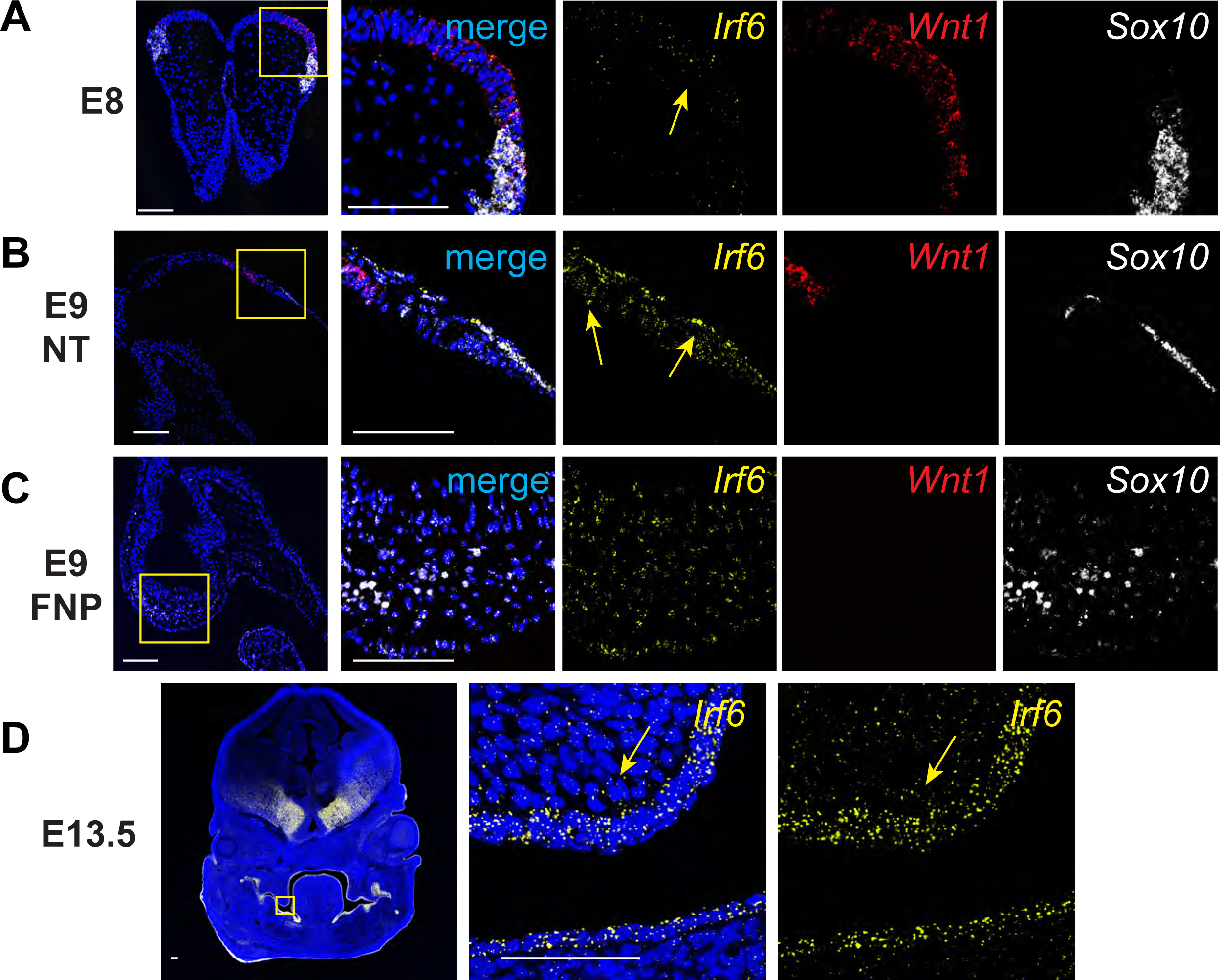
*Irf6* is expressed with neural crest cell markers *Wnt1* and *Sox10* in neural folds and neural tube during early embryogenesis. *In situ* hybridization of *Irf6* (yellow), *Wnt1* (red), and *Sox10* (white) RNA transcripts. **A.** Coronal section of E8 mouse embryo (dorsal to top) showing the neural fold. *In situ* hybridization shows RNA expression domains of *Irf6*, *Wnt1*, and *Sox10*, where *Irf6* and *Wnt1* transcripts are found in the same regions of the neural tube, highlighted by yellow arrow. Box indicates area of higher magnification to the right. **B.** Sagittal section of E9 mouse embryo (cranial to left). Box indicates a magnified portion of the neural tube. *Irf6* is expressed in the neuroectoderm and overlaps with *Wnt1* and *Sox10* expression (yellow arrows). **C.** Sagittal section of E9 mouse embryo (cranial to left). Box indicates a magnified portion of frontonasal prominence (FNP). *Irf6* is expressed in the FNP mesenchyme, along with the migratory NCC marker *Sox10*. **D.** Coronal section of E13.5 embryo (dorsal to top). Box indicates higher magnification of palate shelf epithelium and mesenchyme. *Irf6* is highly expressed in the basal epithelium and periderm and the palate mesenchyme (yellow arrow). Blue is dapi. Scale: 100 uM.

Further, *Irf6* mRNA is expressed within the neural crest-derived craniofacial mesenchyme at E9 (Fig 1c) and E13.5 (Fig 1d). Based on these detailed gene expression findings, as well as previously reported expression of *Irf6* in the neuroectoderm (Bertol et al., 2022; Kousa et al., 2019), we posited that *Irf6* contributes to craniofacial development beyond its established role in the surface epithelium and periderm.

### Generation and validation of an *Irf6* conditional allele

Severe epithelial adhesions and perinatal lethality in the *Irf6* complete knockout embryos (Ingraham et al., 2006) and the *Irf6* R84C single nucleotide substitution mouse (Richardson et al., 2006) impeded full analysis of *Irf6* function. The previously reported *Irf6* floxed mouse allele was reported to exhibit inconsistent recombination depending on the Cre driver used, confounding analysis of *Irf6* requirement in the multiple tissue types (Smith et al., 2017). Given the complexity of *Irf6* gene expression in neuroectoderm and neural crest during early embryogenesis, we generated a new *Irf6* floxed allele to better understand *Irf6* function. We utilized an CRISPReasi technique (Miura et al., 2018) to insert loxP sequences flanking exon 3 and 4 of the *Irf6* gene (Fig 2a). Insertion of the (22bp) loxP sequence was verified by PCR genotyping of potential founders (Fig 2b), followed by Sanger sequencing to confirm loxP insertion without disruption of exonic sequences.

**Fig. 2.**
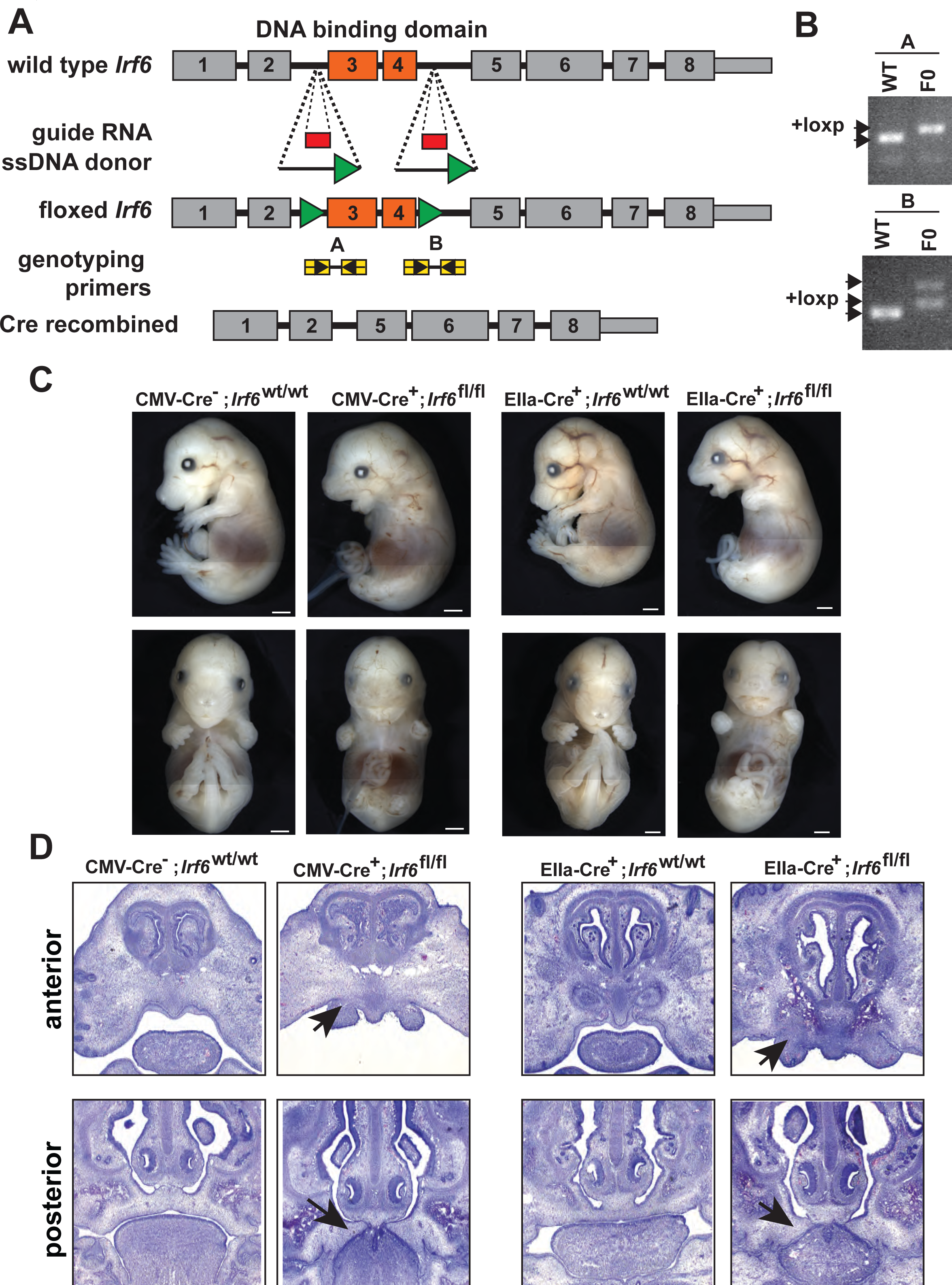
Generation and validation of a conditional *Irf6* null mouse model. **A.** Schematic of gene targeting strategy. Introns flanking *Irf6* exons 3 and 4 were targeted for CRISPR-Cas9-directed homologous recombination with each donor ssDNA containing loxP sequences (green triangles). Insertion of loxP sites into *Irf6* was confirmed by PCR. **B**. and Sanger sequencing. **C.** Cre-mediated recombination was validated using the ubiquitous Cre expressing lines *CMV*-Cre and *EIIa*-Cre. *CMV*-Cre^+^;*Irf6*^fl/fl^ and *EIIa*-Cre^+^; *Irf6*^fl/fl^ mice phenocopied the *Irf6* global KO while Cre*^-^;Irf6*^fl/fl^ and Cre^+^;*Irf6*^wt/wt^ littermates were normal. **D.** Hematoxylin and Eosin staining of coronal sections of E15 *CMV*-Cre or *EIIa*-Cre knockout embryos and littermate controls. Top row is a relatively anterior section while the bottom row is relatively posterior. *CMV*-Cre and *EIIa*-Cre *Irf6* KO embryos phenocopy the dysmorphic alveolar bone and the cleft palate with oral adhesions of the total *Irf6* knockout mouse (arrows).

To test whether Cre expression resulted in recombination and loss of function, the confirmed founder mouse was bred to two different deleter strains; *CMV*-Cre and *EIIa*-Cre. We found that pups that were homozygous for loxP but negative for Cre were phenotypically normal and healthy. Pups that were homozygous for loxP and positive for *CMV*-Cre or *EIIa*-Cre recapitulated the epithelial adhesions, limb abnormalities, and cleft palate displayed by *Irf6* total knockout mice (Fig 2c,d). Further, we found this phenotype to be completely penetrant. Based on these results, we determined faithful recombination of the *Irf6* floxed allele leading to functional Irf6 ablation.

### Ablation of *Irf6* in the *Wnt1* lineage leads to a cranial defect and increased perinatal lethality

With recombination of the floxed *Irf6* allele validated, we tested the effect of *Irf6* ablation in the NCC lineage. We utilized the *Wnt1*-Cre2 and the *Sox10*-Cre mouse lines to drive the recombination of the floxed genome sequence. *Wnt*1 is expressed in the neural folds and pre-migratory NCCs (Lewis et al., 2013; Schock et al., 2017) whereas *Sox10* is expressed in migratory NCCs (Matsuoka et al., 2005). Analysis of *Sox10*-Cre *Irf6* cKO pups revealed no phenotypic effect of *Irf6* ablation in migratory and post-migratory NCCs (data not shown). This finding suggests that although *Irf6* mRNA can be found in NCC-derived mesenchymal tissue, its expression is not necessary for craniofacial development. In contrast to *Sox10*-Cre driven ablation, analysis of *Wnt1*-Cre2 *Irf6* cKO pups revealed a range of phenotype severity with some pups phenotypically normal and viable. We also observed P0 pups that were largely normal but exhibited skin lesions overlying the nasal and frontal bones (Fig 3a). These skin lesions resolved but led to delayed fur growth (Fig 3b). To detect whether *Wnt1*-Cre2 cKO of *Irf6* resulted in differences in pup survival, litter size at P0 was recorded and compared to genotype ratios at 3 weeks of age. Based on total pup numbers and expected ratio (based on parent genotypes), we expected approximately 6 *Wnt1*-Cre2 *Irf6* cKO at weaning. Instead, 1 *Irf6* cKO pup was identified at 3 weeks of age, suggesting perinatal lethality between birth and weaning. The numbers of wild-type and heterozygous pups were as expected. We did not find the lethality of the *Wnt1*-Cre2 *Irf6* cKO pups to be due to cleft palate, as histological examination of P0 dead or moribund pups showed palatogenesis to be normal (Fig. 3c).

**Fig. 3.**
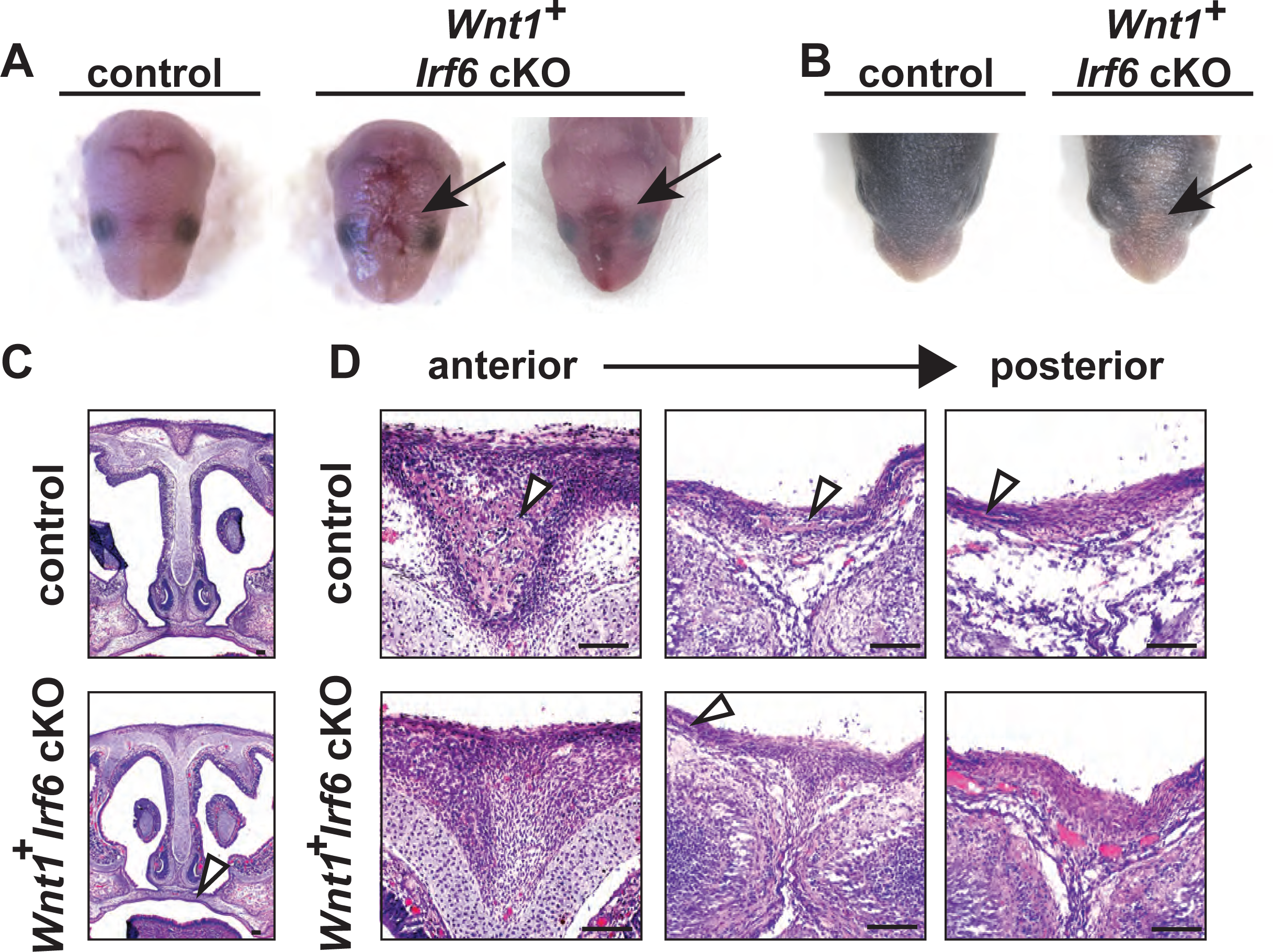
*Wnt1*-Cre-dependent *Irf6* ablation causes cranial defects. **A.** Representative images of littermate control and *Wnt1*-Cre, *Irf6* cKO pups at P0. At parturition, *Wnt1*-Cre^+^;*Irf6*^fl/fl^ cKO mice display midline lesions of varying penetrance (arrow). **B.** Representative images of littermate control and *Wnt1*-Cre^+^;*Irf6*^fl/fl^ cKO pups at P6. As the mouse neonate develops, these frontal lesions resolve but remain evident with deficient or delayed fur growth (arrow). **C.** Hematoxylin and eosin staining of coronal sections through the palate of E16 *Wnt1*-Cre^+^;*Irf6*^fl/fl^ cKO and littermate control embryos shows normal development (arrow). **D.** Hematoxylin and eosin staining of coronal sections through the nasal and frontal bones of *Wnt1*-Cre^+^;*Irf6*^fl/fl^ cKO and littermate control. Sections move anterior to posterior from left to right. Bone tissue is indicated with arrows. *Wnt1*-Cre^+^;*Irf6*^fl/fl^ cKO mice have a lack of cranial bone development and suture formation at the midline (bone tissue indicated by arrows). Scale: 100 µM.

To examine whether the underlying calvarial development was affected in the *Wnt1*-Cre2 *Irf6* cKO before parturition, we performed histology on coronal sections taken through the nasal-frontal bone junction of *Wnt1*-Cre2 *Irf6* cKO E16 pups and littermate controls. We found that control mice had bone tissue at the midline, forming a suture between right and left calvaria. In contrast, the *Wnt1*-Cre2^+^;Irf6^fl/fl^ cKO mice exhibited a large gap devoid of bone tissue that spanned the midline (Fig 3d). These findings of a midline cranial defect and partial lethality in *Wnt1*-Cre2 *Irf6* cKO mice suggest that *Irf6* expression in the pre-migratory NCCs is functionally required for craniofacial development.

### *Wnt1*-Cre2 *Irf6* cKO mice exhibited incomplete frontal and parietal bone development

The variable severity of the cranial defect in *Wnt1*-Cre *Irf6* cKO mice spurred us to examine the cranial bone development of the mice more precisely with microCT analysis. *Wnt1*-Cre cKO pups and sex-matched littermate controls were collected at 10 days of age for microCT scanning. For controls, we analyzed both *Wnt1*-Cre negative, *Irf6*^fl/fl^ and *Wnt1*-Cre positive, *Irf6*^wt/fl^ pups to account for potential differences caused by the *Wnt1*-Cre transgene. *Wnt1-* Cre*^+^;Irf6^fl/fl^* pups exhibited decreased mineralization of the frontal and parietal bones at the midline, although the degree of this defect was variable between individuals (Fig 4a). These observations are similar to previously published microCT analysis of the *Irf6* null mouse (Thompson et al., 2019). To quantify potential changes in cranial development in the cKO mice, we performed a series of measurements based on established anatomical landmarks in cKO versus sex-matched littermate *Wnt1*-Cre negative controls (Fig 4a). Overall, we did not detect significant differences in length or width measurements of the frontal, maxillary, or nasal bones, except that the width across the anterior portion of the frontal bones of the *Wnt1*-Cre2^+^;*Irf6*^fl/fl^ cKO mice was significantly larger (Fig. 4b). Nasal bones of *Wnt1*-Cre2^+^;*Irf6*^fl/fl^ cKO mice tended to be shorter, however, this measurement was variable in the control pups (Fig. 4b). These data demonstrate that *Irf6* expressed in NCCs contributes to a midline calvarial bone defect. We hypothesize this to be an indirect effect of loss of *Irf6* in the neural folds, as non-NCC derived calvarial tissue, namely parietal bone, is also deficient in the *Wnt1*-Cre *Irf6* cKO mice.

**Fig. 4.**
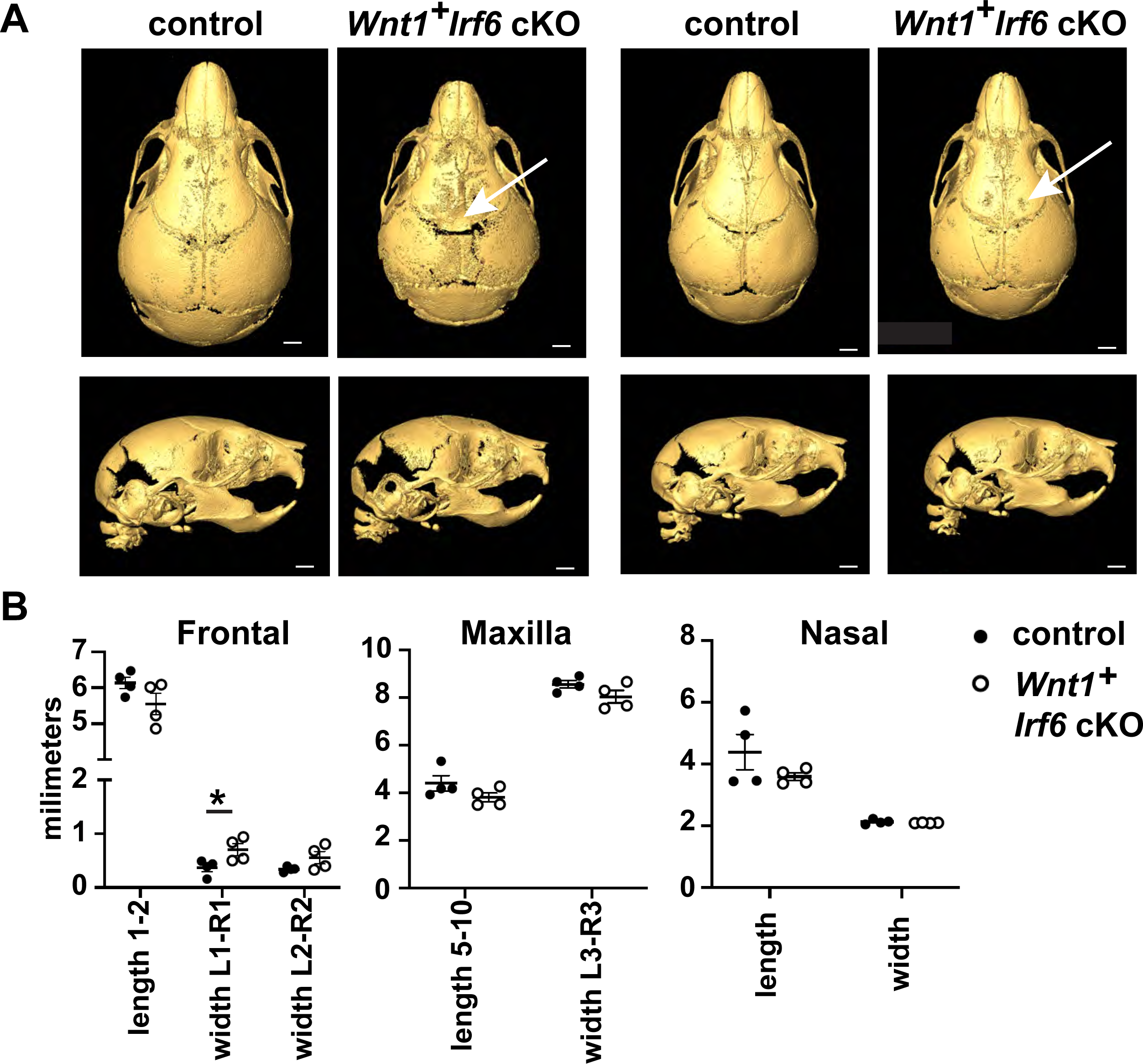
Cranial bone development is impaired in *Wnt1-*Cre *Irf6* cKO mice. **A.** Representative microCT reconstructions of P10 *Wnt1*-Cre^+^;*Irf6*^fl/fl^ cKO mice and littermate sex-matched controls. *Wnt1*-Cre^+^;*Irf6*^fl/fl^ cKO mice have decreased formation or mineralization of the cranial bones at the midline with variable penetrance (arrows). Scale: 1 mm. **B.** MicroCT reconstructions were utilized for cranial bone measurements. The space between the left and right frontal bones of *Wnt1*-Cre^+^;*Irf6*^fl/fl^ cKO mice was significantly wider than controls (L1-R1, *p<0.05) and the frontal bones tended to have decreased total length (length 1-2). Maxilla of *Wnt1*-Cre^+^;*Irf6*^fl/fl^ cKO mice tended to be smaller (lower length and width measurements) and the frontal bone of *Wnt1*-Cre^+^;*Irf6*^fl/fl^ cKO mice tended to be shorter, however, these differences were not significantly different. N=4.

### *Wnt1*-Cre dependent *Irf6* ablation altered neuroepithelial morphology and *Wnt1* **expression**

The cranial defect observed in *Wnt1*-Cre;*Irf6* cKO mice involved both the overlying skin (Fig 3a) and the underlying cranial bone (Fig 3d). Further, neural crest-derived frontal bone and non-neural crest-derived parietal bone were affected by *Wnt1*-Cre;*Irf6* cKO (Fig. 4a). Therefore, we reasoned that this phenotype may be the manifestation of a rostral neural tube closure defect. We examined the neural folds of E8 *Wnt1*-Cre^+^;*Irf6*^fl/f^ cKO and littermate control embryos. Transverse sections through the cranial neural folds showed differences in overall morphology, with the cKO embryos tending to be more elongated anterior-posterior as compared to controls (Fig. 5). Further, *Wnt1* expression in the *Wnt1*-Cre^+^;*Irf6*^fl/fl^ cKO embryos was laterally displaced relative to the more posterior expression observed in the controls (Fig. 5). These results are consistent with a previous report of dysmorphic neuroectoderm and neural fold morphology in *Irf6* null embryos (Bertol et al., 2022).

**Fig. 5.**
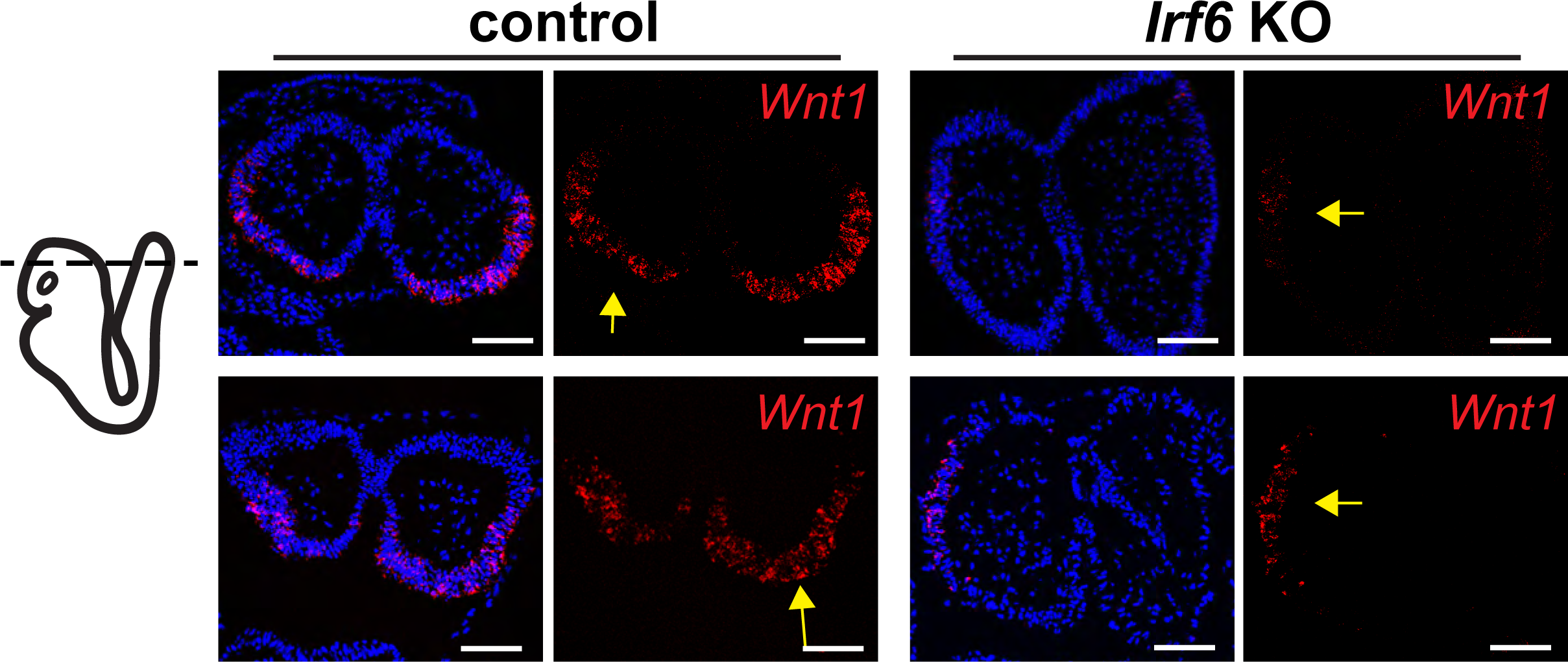
*Irf6* ablation in the neuroectoderm and neural crest changes *Wnt1* expression domains within the neural folds. **A.** RNAscope *in situ* hybridization of transverse sections of *Wnt1*-Cre^+^;*Irf6*^fl/fl^ cKO and littermate control E8 embryos. Rows represent 2 individuals of each genotype. Whereas *Wnt1* expression (red) is localized to the caudal-dorsal neural folds in the control embryos, *Wnt1* expression in *Wnt1*-Cre^+^;*Irf6*^fl/fl^ cKO embryos is displaced laterally (arrows). Blue is dapi. Scale: 100 µM

### Ablation of *Irf6* in the periderm causes a milder global disruption phenotype

To examine the phenotypic effects of *Irf6* ablation in the periderm, we utilized the *Krt6ai*-Cre driver line where the keratin 6 promoter drives Cre expression predominantly in the oral periderm after E14.5 (Saroya et al., 2023). *Krt6ai*-Cre^+/-^;*Irf6*^wt/fl^ males were bred to *Irf6*^fl/fl^ females and it was noted that the *Krt6ai*-Cre^+^;*Irf6*^fl/fl^ genotype was not found at 3 weeks of age. As *Irf6* global null mice die shortly after birth, we observed neonates at P0 and found that a few lacked a milk spot and appeared to be failing to thrive. Pups were collected and genotyping determined these unhealthy pups to be *Krt6ai*-Cre^+^;*Irf6*f^l/fl^ whereas healthy pups were negative for Cre or were *Irf6*^wt/fl^. Closer examination of P0 neonates revealed shiny skin as has been previously noted for KO (Ingraham et al., 2006). *Krt6ai-*Cre^+^;*Irf6*f^l/fl^ pups also exhibited pterygium of the fore and hind limbs consistent with a milder form of the cocooning observed in the global *Irf6* null mouse (Ingraham et al., 2006) (Fig. 6A,B). *Krt6ai*:Cre^+^;*Irf6*^fl/fl^ neonates exhibited simple syndactyly digits of the fore and hind limbs (Fig. 6B).

**Fig. 6.**
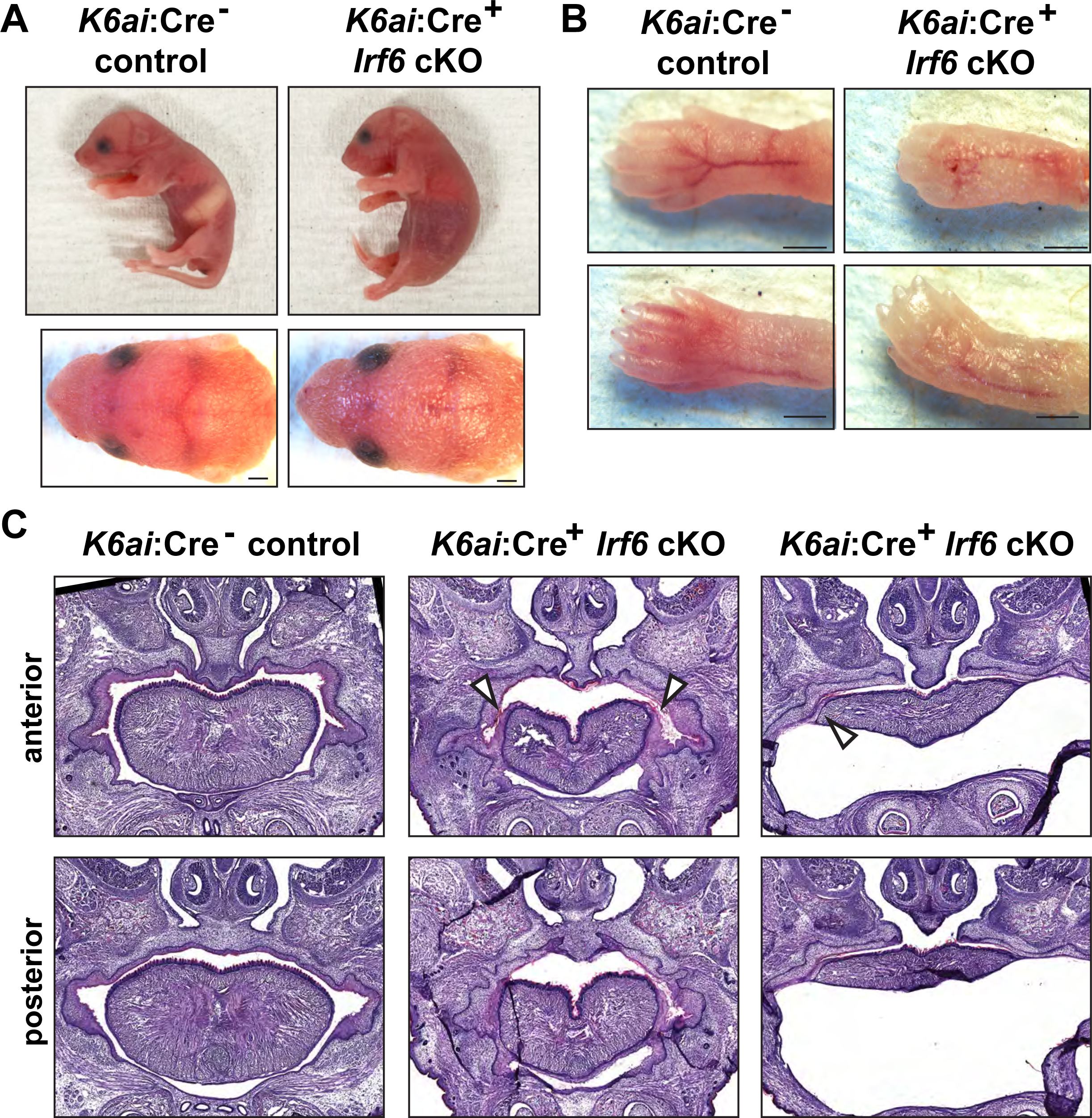
Periderm-specific ablation of *Irf6* results in a comparable but mild form of the global *Irf6* KO phenotype. *Krt6ai*-Cre^+^;*Irf6*^fl/fl^ and littermate control neonates were collected at P1. **A.** Lateral and caudal representation of neonates comparing control *Krt6ai*-Cre^-^;*Irf6*^fl/fl^ with *Krt6ai*-Cre^+^;*Irf6*^fl/fl^ cKO. **B.** *Krt6ai*-Cre^-^;*Irf6*^fl/fl^ exhibit normal skin and digits; however *Krt6ai*-Cre^+^;*Irf6*^fl/fl^ reveal abnormal skin and fused digits phenotype. Scale: 500 µM. **C.** Hematoxylin and Eosin staining of coronal sections through vomeronasal and primary palate of neonates. *Krt6ai*-Cre^-^;*Irf6*^fl/fl^ mice show normal septum and palate. *Krt6ai*-Cre^+^;*Irf6*^fl/fl^ mice reveal abnormal septum and adhesions of the tongue.

The lack of a milk spot in the *Krt6ai*-Cre^+^;*Irf6*f^l/fl^ neonates suggested impaired feeding and possible palate defects and oral adhesions as occur with global *Irf6* ablation. Histological examination revealed that *Krt6ai*-Cre^+^;*Irf6*^fl/fl^ mice present with lateral adhesions of the tongue to the oral cavity and a cleft of the secondary palate of variable penetrance (Fig. 6C). In some *Krt6ai*-Cre^+^;*Irf6*^fl/fl^ individuals we found sublingual fluid accumulation that we presume to be caused by the oral adhesions. No differences were observed in the lip or primary palate.

To compare *Krt6ai* periderm-specific *Irf6* ablation findings to pan-epithelial ablation, we utilized the *Crect* driver line. The *Crect* mouse has been previously utilized to conditionally ablate gene expression in the ectoderm, including the oral and cranial epithelium (Reid et al., 2011; Schock et al., 2017). *Crect*^+^;*Irf6*^fl/fl^ embryos were examined at approximately E17 and were found to recapitulate the *Irf6* knockout phenotype with abnormal skin, foreshortened limbs, and deficient development of the maxilla and mandible (Fig. S1). Histology of these mice showed adhesion of the tongue to the palate, similar to *Irf6* global null mice (Fig. S1). This finding suggests that *Crect* expression largely overlaps with the expression of endogenous *Irf6* gene expression, leading to complete *Irf6* ablation in the *Crect*^+^;*Irf6*^fl/fl^ cKO mouse.

## Discussion

Mutations in *IRF6* underlie VWS and PPS, which are characterized by varying degrees of cleft lip, cleft palate, lip pits, skin folds, syndactyly, and oral adhesions (REF). *Irf6* null and the *Irf6*R84C mutant mouse models recapitulate aspects of these syndromes with severe oral adhesions, surface epithelium adhesions, and dysfunctional keratinocytes which cause neonatal lethality (Ingraham et al., 2006; Kondo et al., 2002). *IRF6* is also associated with non-syndromic cleft lip and palate (Leslie et al., 2016), and yet the severe adhesions of the tongue within the oral cavity in the *Irf6* null and *Irf6R84C* mutant mouse models complicate a direct comparison to the human condition. This study generated a new *Irf6* conditional knockout mouse model and demonstrated reliable recombination of the conditional allele when tested with various Cre driver lines. This new conditional Irf6 allele facilitated the investigation of tissue-specific roles of *Irf6*.

*IRF6*, *TFAP2A*, and *GRHL3* share a genetic regulatory pathway and ablation of each of these genes in mice causes similar cleft, skin, and limb defects (Ingraham et al., 2006; Kousa et al., 2019; Richardson et al., 2006; Schorle et al., 1996; Siewert et al., 2023; Smith et al., 2017; Ting et al., 2003; Zhang et al., 1996). As such, it is intriguing that Tfap2a and Grhl3 are associated with neural tube defects, whereas defects are not observed in the Irf6 ablated mice (Schorle et al., 1996; Ting et al., 2003; Zhang et al., 1996). To investigate this phenomenon Kousa et al., developed an *Irf6* loss-of-function and gain-of-function allelic series in mice and found rostral neural tube defects associated with *Irf6* overexpression and caudal defects associated with *Irf6* loss of function (Kousa et al., 2019). We hypothesized that the severe epithelial adhesions resulting from periderm dysfunction in the *Irf6* null mouse may mask neural tube defects and we therefore generated a conditional KO where *Irf6* would be ablated in *Wnt1* expressing neuroectoderm and neural crest cells, including those in the neural folds. We found a rostromedial defect in these mice of varying severity that affected the skin and calvarial bone. Further, we found changes to neural fold morphology and *Wnt1* expression patterns in these embryos. Together, these data corroborate a role for *Irf6* in the patterning and morphogenesis of the rostral neural tube in mice. Differences in phenotype and severity between our results and Kousa et al. may be attributed to spatial and temporal differences in the respective overexpression and knockout drivers that were utilized (*Krt14* versus *Wnt1*). Further, additional neural tube phenotypes may become apparent in the *Wnt1*-Cre *Irf6* cKO upon combinatorial genetic disruption of *Tfap2a* or *Grhl3*.

*Irf6* is widely expressed in the pan-epithelium and its specific role in various epithelial populations (i.e. basal epithelium versus periderm) and those contributions to the mutant phenotype have had limited direct investigation. Kousa et al. previously investigated the role of *Irf6* in the basal epithelium by utilizing the *Krt14* promoter to express *Irf6* in the basal epithelium on an *Irf6* global null background. It was found that *Irf6* expression in the basal epithelium partially rescued some aspects of the *Irf6* null phenotype, namely the skin adhesions of the axial and appendicular skeleton but did not rescue the cleft palate (Kousa et al., 2017). Utilizing our *Irf6* floxed mouse and the *Krt6ai*-Cre driver, we found that ablation of *Irf6* in the periderm largely phenocopied the *Krt14:Irf6^tg^* rescue. Limb defects were similar in that the limbs were not adhered to the body yet syndactyly of the digits were observed. Whereas Kousa et al. reported oral adhesions slightly less severe than the global KO and cleft palate, the periderm-specific *Irf6* KO mice had relatively mild oral adhesion and cleft of the palate was incompletely penetrant. Therefore, our data coincide with previous findings, and differences in phenotype and severity are likely due to differences in cell specificity and timing of expression.

Irf6 has a key role in the regulation of epithelial proliferation and differentiation (Bailey et al., 2008; Biggs et al., 2012; Girousi et al., 2021; Oberbeck et al., 2019). As such, *IRF6* is implicated in epidermal wound healing and children with VWS have an increased risk of wound complications following surgical repair of orofacial clefts (Hixon et al., 2017; Jones et al., 2010; Rhea et al., 2020). Further, loss of Irf6 expression is associated with epidermal malignancy (Botti et al., 2011; Darido et al., 2016; Parisi et al., 2022; Yan et al., 2023). Investigation into these roles of Irf6 have, until now, depended on human patient-derived cells, genetically manipulated cell lines, and gene association studies. The availability of this *Irf6* conditional mouse allele will allow post-natal ablation of *Irf6* and facilitate mechanistic studies of epithelial biology in a mouse model.

This study successfully generated and validated a conditional *Irf6* mouse allele. This mouse model will serve as an invaluable tool for advancing our comprehension of *Irf6*’s multifaceted functions and for developing targeted interventions for conditions like orofacial clefts, wound healing complications, and various cancers.

## Acknowledgments

CRISPR design consultation, zygote microinjection, and embryo implantation were performed by The Genome Modification Facility and Harvard University.

MicroCT scanning was performed by the Center for Musculoskeletal Research Imaging and Biomechanical Testing Core (NIH P30 AR070542).

---

**Fig. S1.**
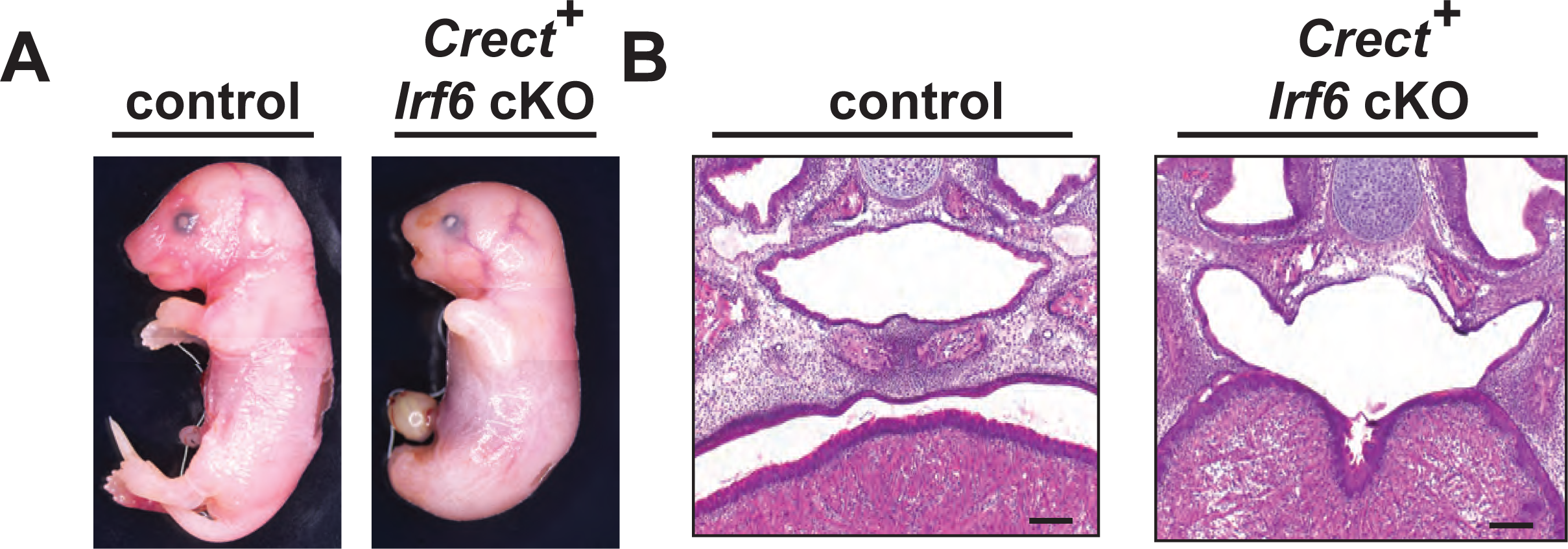
*Crect*-driven *Irf6* ablation recapitulates the global *Irf6* KO phenotype. **A.** Representative images of littermate control and *Crect*^+^-*Irf6^fl/fl^* cKO pups at approximately E17. *Crect*^+^-*Irf6^fl/fl^* pups exhibit “cocooning” taught skin, abnormal and shortened limbs, and an umbilical hernia that has been described for the *Irf6* global KO. **B.** Hematoxylin and eosin staining of coronal sections of approximately E17 *Crect*^+^-*Irf6^fl/fl^* pup and littermate control. *Crect*^+^-*Irf6^fl/fl^* cKO pups exhibit severe oral adhesions and cleft palate similar to the global *Irf6* KO mouse. Scale: 100 µM.

